# Metadichol, a modulator that controls expression of Toll Like Receptors in Cancer Cell Lines

**DOI:** 10.1101/2024.08.18.608483

**Authors:** P.R. Raghavan

## Abstract

Toll-like receptors are essential players in the innate immune signaling system and play critical roles in recognizing pathogen-associated and damage-associated molecular patterns. Its presence in various cancer cells, including breast, ovarian, cervical, colon, hepatocellular carcinoma, prostate, lung cancer, melanoma, and neuroblastoma cells, has been associated with cancer progression, immune evasion, apoptosis, and chemoresistance. Despite extensive research, no small molecules can induce the expression of all TLRs in cancer cells have been identified. This study investigated the effects of metadichol, a nanoemulsion of long-chain alcohols, on the expression of TLR receptor families 1–10 in cancer cell lines. Using quantitative polymerase chain reaction (qPCR) and other molecular biology techniques, we assessed the concentration-dependent effects of metadichol on TLR expression, which ranged from 1 picogram per ml to 100 nanograms per ml. Our results show that metadichol can modulate TLR expression in a cancer cell type-specific manner, with TLR4 being upregulated in lung cancer cells, enhancing antitumor effects but downregulated in other cancer lines, reducing inflammation.

Furthermore, TLR7, TLR8, and TLR9 are upregulated in certain cell lines. Importantly, metadichol induced the downregulation of MYD88, TRAF6, and IRAK4 across various cell lines, suggesting considerable potential to inhibit tumor growth, enhance apoptosis, and reduce metastasis. This broad-spectrum gene activation supports a multitarget therapeutic approach, making metadichol the first molecule to target the entire TLR family without the need for complex biological methods such as CRISPR—alternatively, virus grafting techniques. Our findings underscore the potential of metadichol in treating complex diseases involving multiple pathways, such as cancer, chronic inflammatory pathways, and infectious diseases, highlighting noteworthy advancements in cancer therapy.

## Introduction

Toll-like receptors (TLRs) are a family of pattern recognition receptors that play essential roles in the innate immune response and recognize pathogen-associated molecular patterns (PAMPs) and damage-associated molecular patterns (DAMPs). (1) We recently reported (2) the expression of TLR1-10 and MYD88, TICAm1, IRAJ4, Traf3, and TRAF6 in PBMCs at concentrations ranging from 1 picogram to 100 nanograms. All 15 genes were expressed concentration-dependent and exhibited an inverted U response, except for TLR4, whose expression was downregulated. The expression of TLRs in cancer cells has been extensively studied, revealing both tumor-promoting and tumor-suppressing roles depending on the context. (3) The engagement of TLRs in cancer cells can have dual effects, either promoting or inhibiting cancer progression, depending on the context and specific TLRs involved.

The specific challenge addressed in this study is the modulation of TLR expression in cancer cell lines by metadichol, a nanoemulsion of long-chain alcohols. TLRs are expressed on immune cells and various cancer cells, influencing cancer progression and the tumor microenvironment. Different types of cancer cells express various TLRs. For example, gastric cancer cells express TLR2, TLR4, TLR5, and TLR9, whereas breast cancer cells express TLR1, TLR2, TLR3, TLR4, TLR5, TLR9, and TLR10. TLR activation in cancer cells can produce cytokines and chemokines, which can recruit immune cells and drive the inflammatory response in the tumor microenvironment. These transcription factors can promote or inhibit tumor growth depending on the context. (4)

Our study investigated metadichol (5), an emulsion of long-chain alcohols, induced expression of TLR receptors (1--10), and five other downfield genes of the TLR network pathway in cancer cell lines, aiming to explore the potential of metadichol as a modulator of TLR expression and its implications for cancer therapy. We used quantitative reverse transcriptase polymerase chain reaction (q-RT PCR) to measure changes in TLR mRNA.

## Materials and methods

All work was outsourced commercially to the service provider Skanda Life Sciences Pvt Ltd., Bangalore, India. The primers used were obtained from Saha BioSciences, Hyderabad, India. The raw qRT PCR data are provided in the Supplemental files.

### 1 ) Cell treatment

The cells were treated for 24 hrs at the following concentrations in growth media without FBS.

**Table 1:**
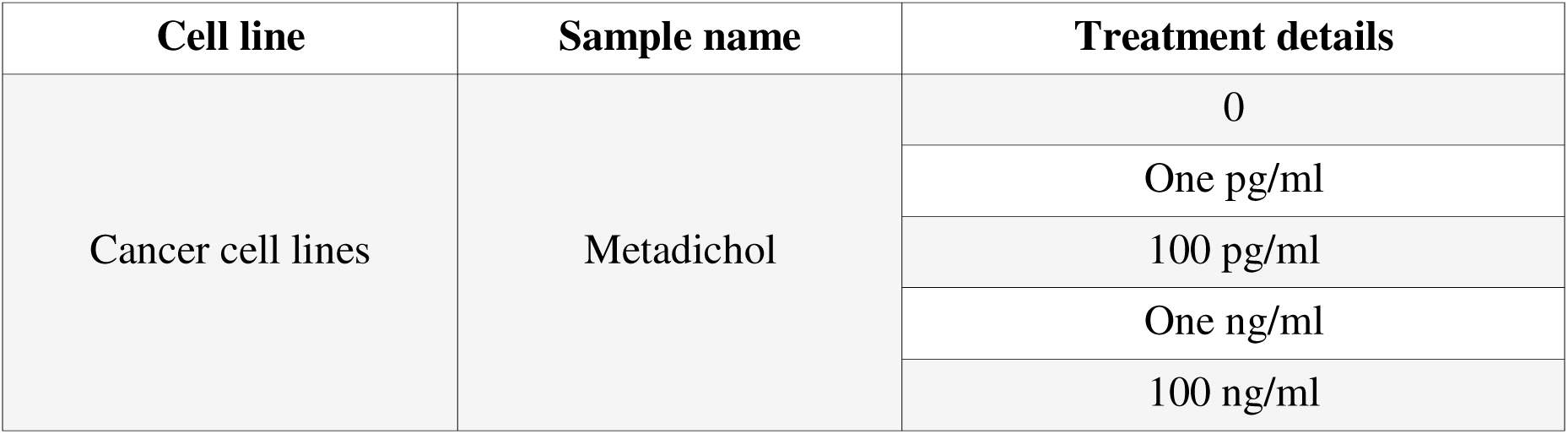
Treatment concentrations.

| Cell line | Sample name | Treatment details |
| --- | --- | --- |
| Cancer cell lines | Metadichol | 0 |
|  |  | One pg/ml |
|  |  | 100 pg/ml |
|  |  | One ng/ml |
|  |  | 100 ng/ml |

## Sample Preparation and RNA Isolation

The treated cells were dissociated, rinsed with sterile 1X PBS, and centrifuged. The supernatant was decanted, and 0.1 ml of TRIzol was added and gently mixed by inversion for 1 min. The samples were allowed to stand for 10 min at room temperature. To this mixture, 0.75 ml of chloroform was added per 0.1 ml of TRIzol used. The contents were vortexed for 15 seconds. The tube was allowed to stand at room temperature for 5 mins. The resulting mixture was centrifuged at 12,000 rpm for 15 min at four °C. The upper aqueous phase was collected in a new sterile microcentrifuge tube, to which 0.25 ml of isopropanol was added, and the mixture was gently mixed by inverting the contents for 30 seconds and then incubated at -20 °C for 20 minutes. The contents were centrifuged at 12,000 rpm for 10 minutes at four °C. The supernatant was discarded, and the RNA pellet was washed by adding 0.25 ml of 70% ethanol. The RNA mixture was subsequently centrifuged at 12,000 rpm at four °C. The supernatant was carefully discarded, and the pellet was air-dried. The RNA pellet was then resuspended in 20 µl of DEPC-treated water. The total RNA yield was quantified via a Spectra drop (Spectramax i3x, Molecular Devices, USA).

**Table 2:** Total RNA yield (shown here as an example Jurkat line)

|  | Test concentrations |  |  |  |  |
| --- | --- | --- | --- | --- | --- |
| RNA yield (ng/µl) | 0 | One pg/ml | 100 pg/ml | One ng/ml | 100 ng/ml |
| JURKAT | 768.8 | 310.25 | 631.28 | 385.6 | 472.169 |

### QRT-RT**□**PCR analysis

#### cDNA synthesis

cDNA was synthesized from 500 ng of RNA via a cDNA synthesis kit from the Prime Script RT reagent kit (TAKARA) with oligo dT primers according to the manufacturer’s instructions. The reaction volume was set to 20 μl, and cDNA synthesis was performed at 50 °C for 30 min, followed by RT inactivation at 85 °C for 5 min using Applied Biosystems, Veritii. The cDNA was further used for real-time PCR analysis.

### Primers and qPCR analysis

The PCR mixture (final volume of 20 µL) contained 1 µL of cDNA, 10 µL of SYBR Green Master Mix, and one µM complementary forward and reverse primers specific for the respective target genes. The reaction conditions were as follows: initial denaturation at 95 °C for 5 min, followed by 30 cycles of secondary denaturation at 95 °C for 30 s, annealing at the optimized temperature for 30 s, and extension at 72 °C for 1 min. The results obtained were analyzed using CFX Maestro software.

The fold change was calculated via the ΔΔCT method.

The comparative CT method was used to determine the relative expression of target genes to that of the housekeeping gene (β-actin) in untreated control cells.

The delta CT for each treatment was calculated via the following formula:

Delta Ct = Ct (target gene) – Ct (reference gene).

To compare the delta Ct of individually treated samples with that of the untreated control samples, the Ct of each group was subtracted from that of the control to obtain the delta delta CT.

Delta delta Ct = delta Ct (treatment group) – delta Ct (control group).

The fold change in target gene expression for each treatment group was calculated via the formula: Fold change = 2^ (−delta Ct).

**Table 3:**
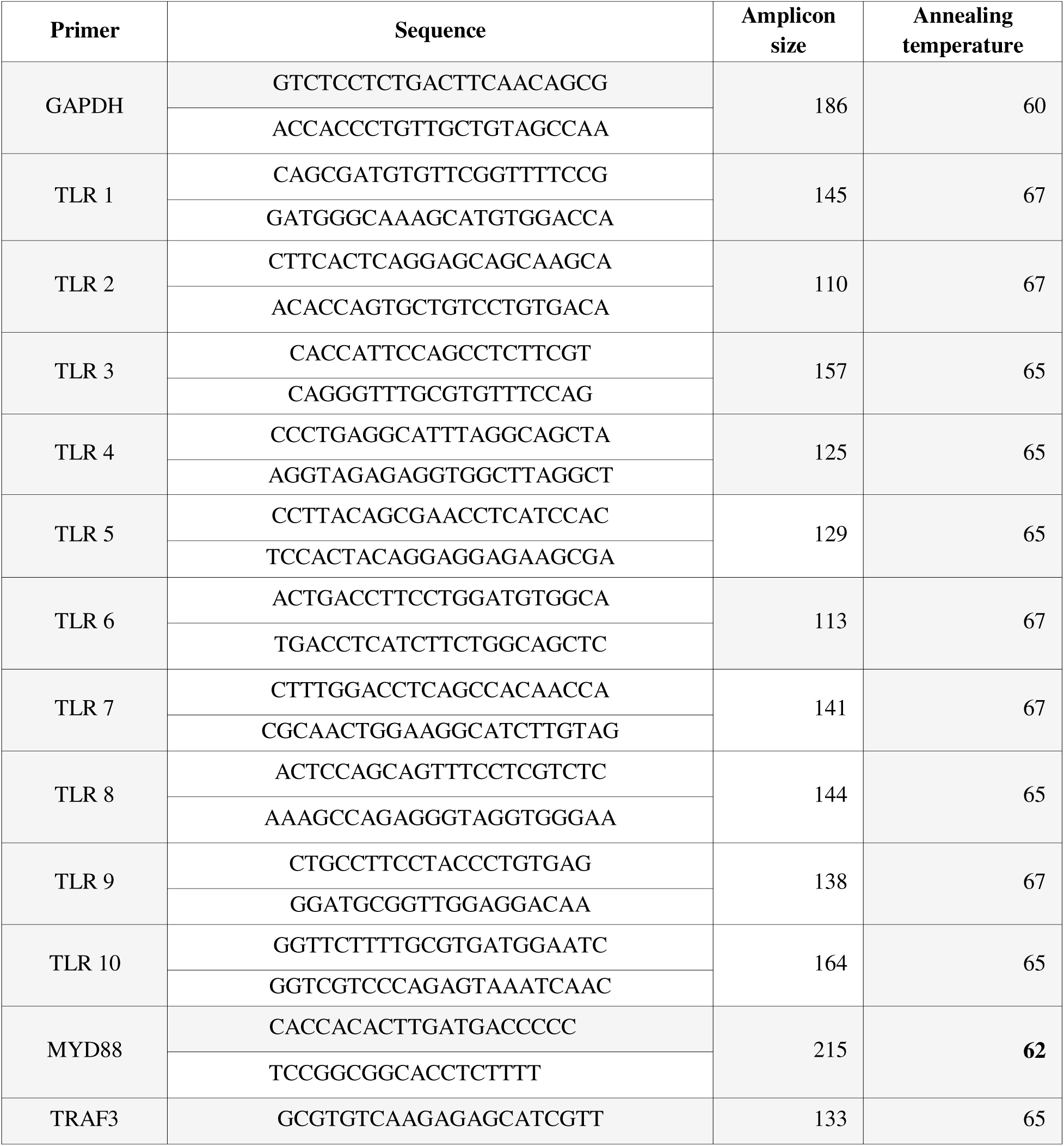
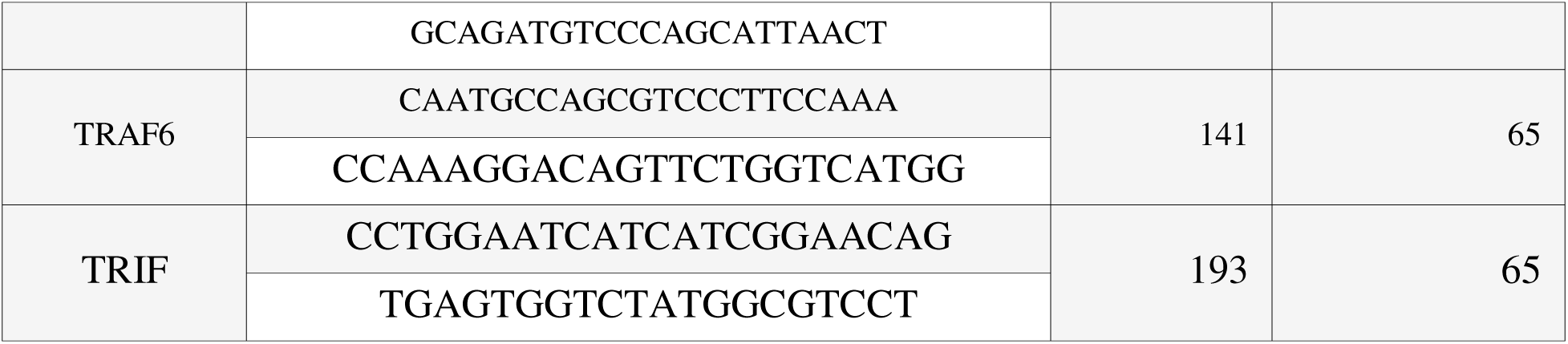
Primer details.

## Results

The results of the TLR expression analysis of the eight cancer lines studied are shown in the figures (TLR1-10, MYD88, TRAF3, TEAF6, IRAK4, and Ticam1) below. For significance, we cut off a 2-fold increase in expression and a 50% reduction in downregulation. Red and gray dashed lines, respectively, represent these. The key findings of this study revealed that Metadichol induces significant changes in the expression levels of TLR1--10 and downfield genes of the TLR network pathway (6) MYD 888, IRAK4, TICAM1, TRIF3, and TRIF6 in cancer cell lines. The gene expression occurred in a dose-dependent manner. One common thread is that at 100 nanograms, the expression of TLRs is inhibited across the board. TLR4 is completely downregulated at all concentrations except in the case of A-549 lung cells, where it is upregulated as TLR4, known to act as an antitumor agent in lung cancer. With a few exceptions @ 100 nanograms, most TLR1-10 and downregulated gene expression are inhibited/downregulated. Overall, the downregulation of TLRs at high concentrations can play an essential role in cancer treatment and inflammation and enhance immune responses against tumors.

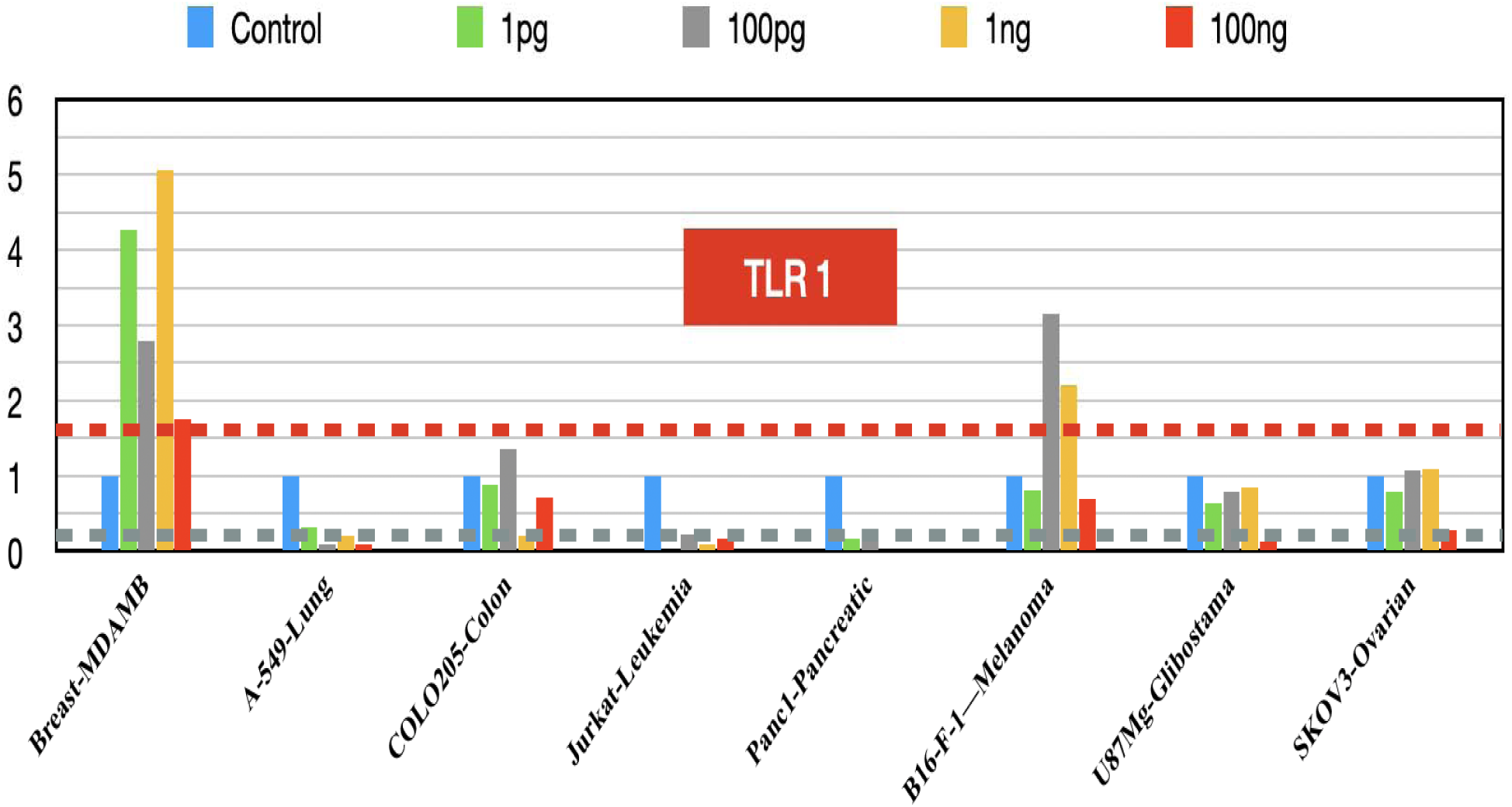

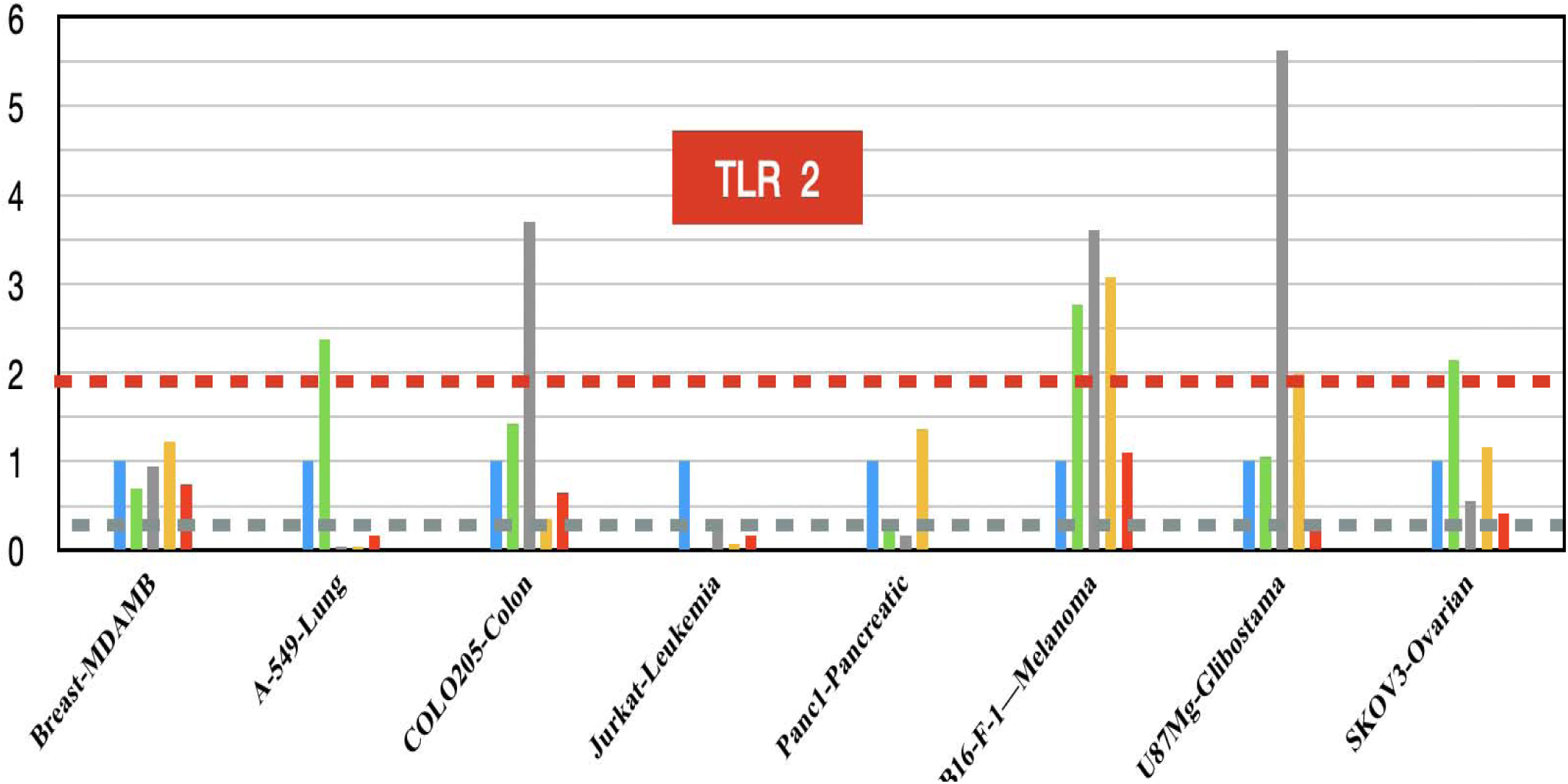

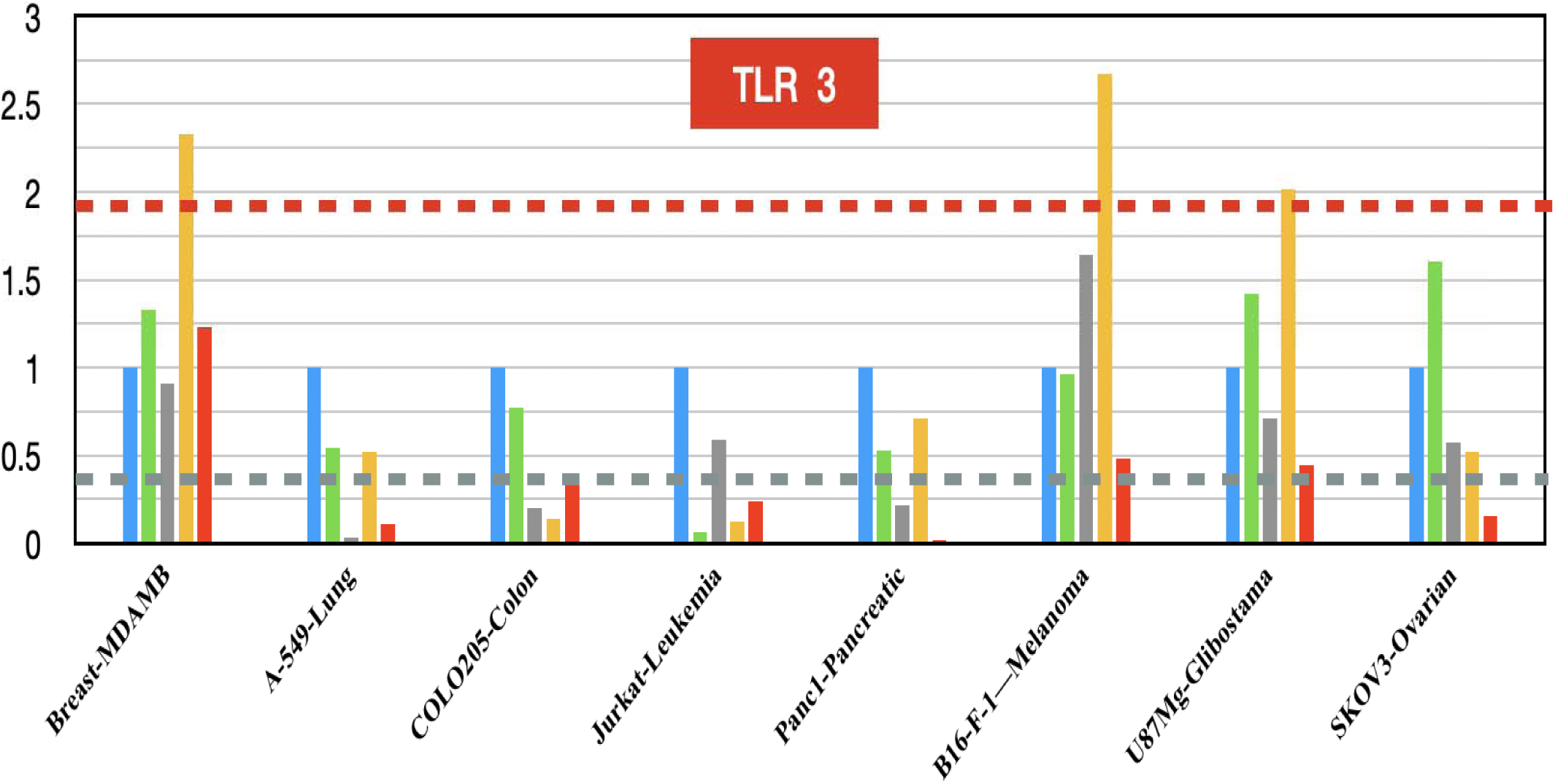

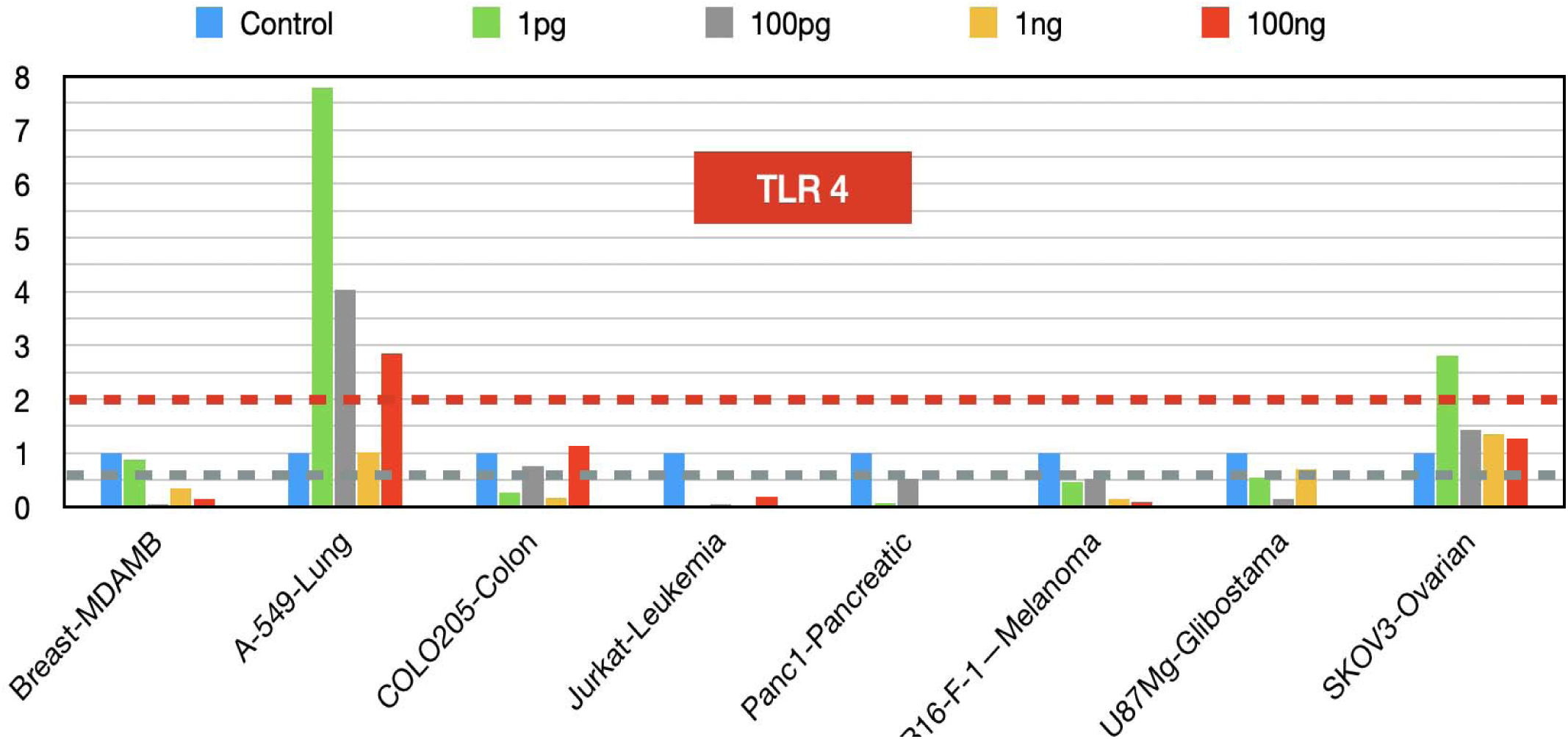

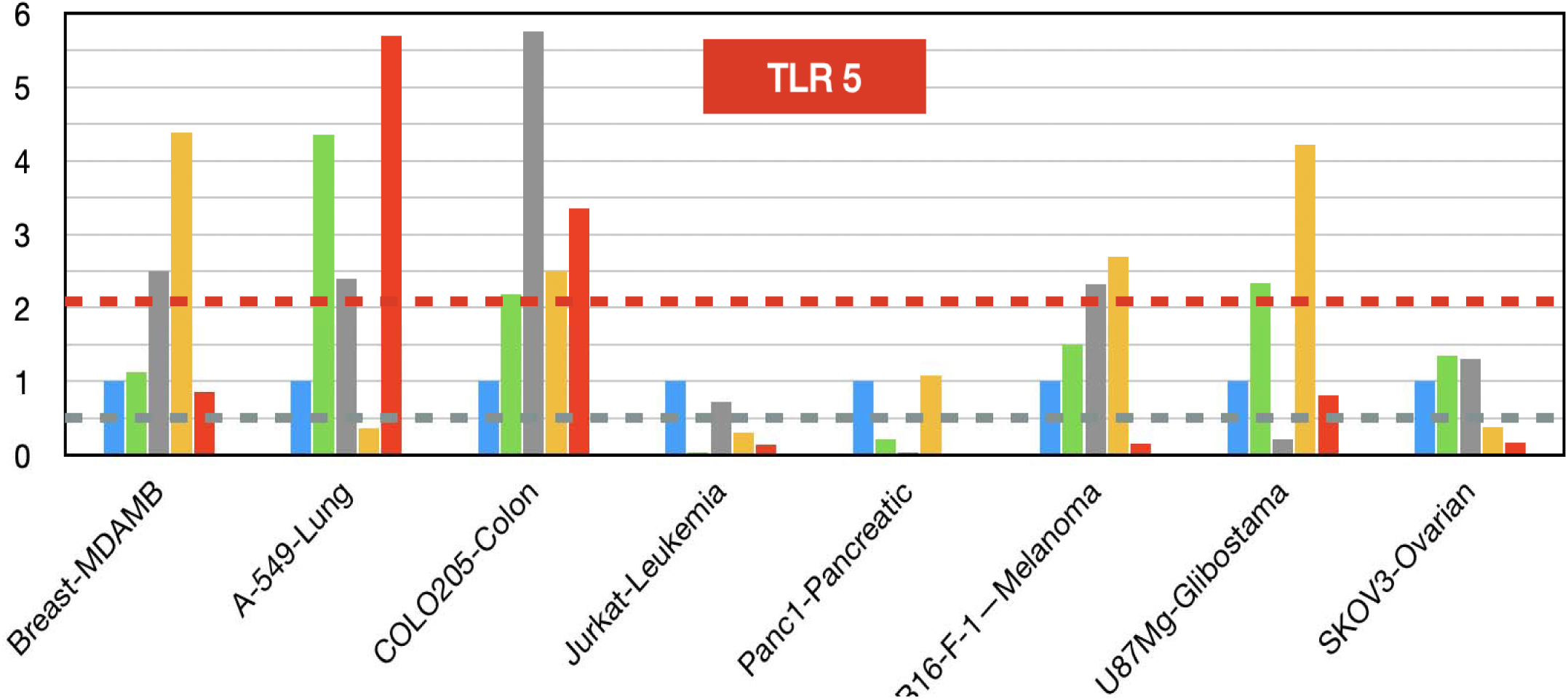

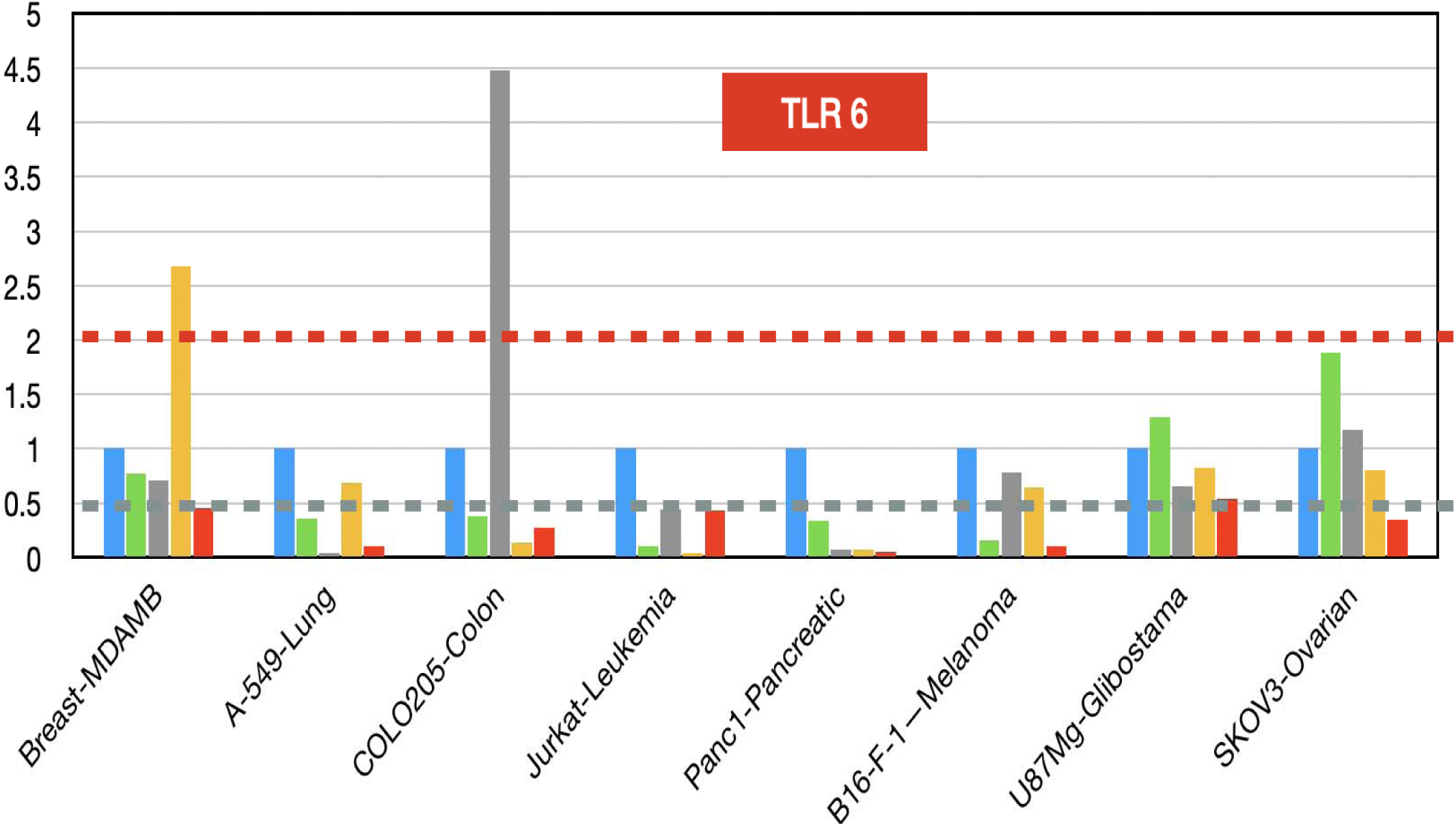

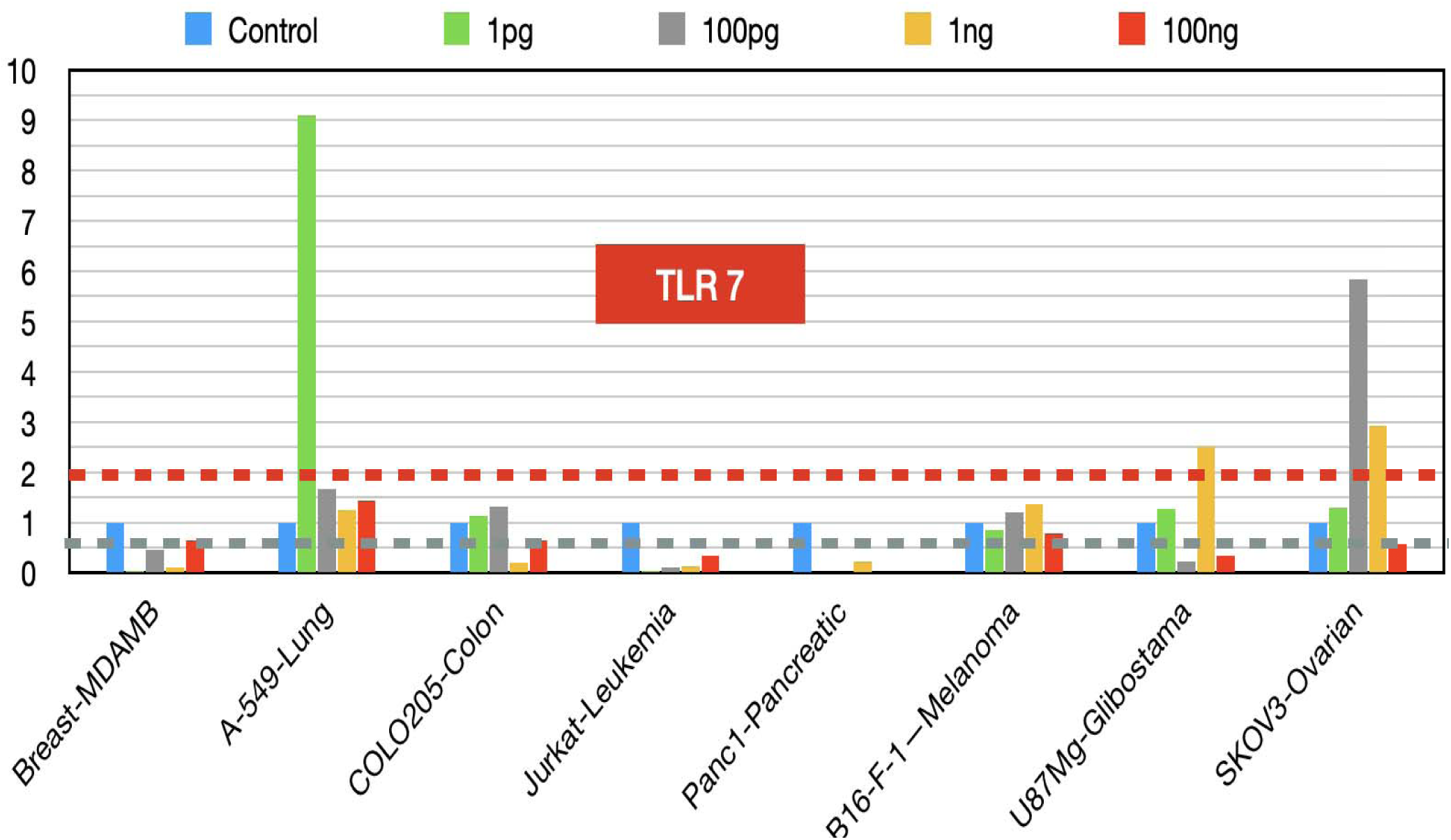

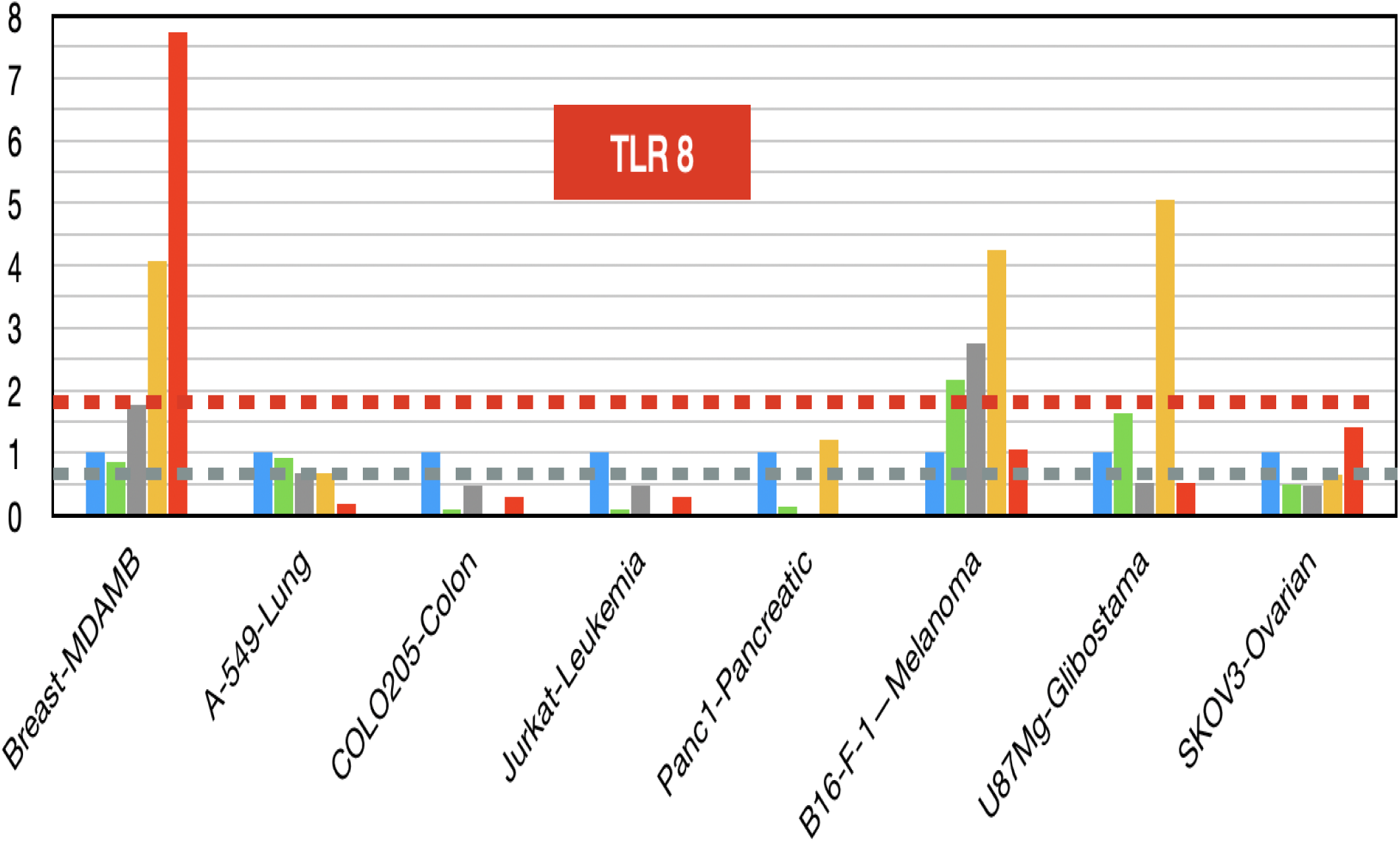

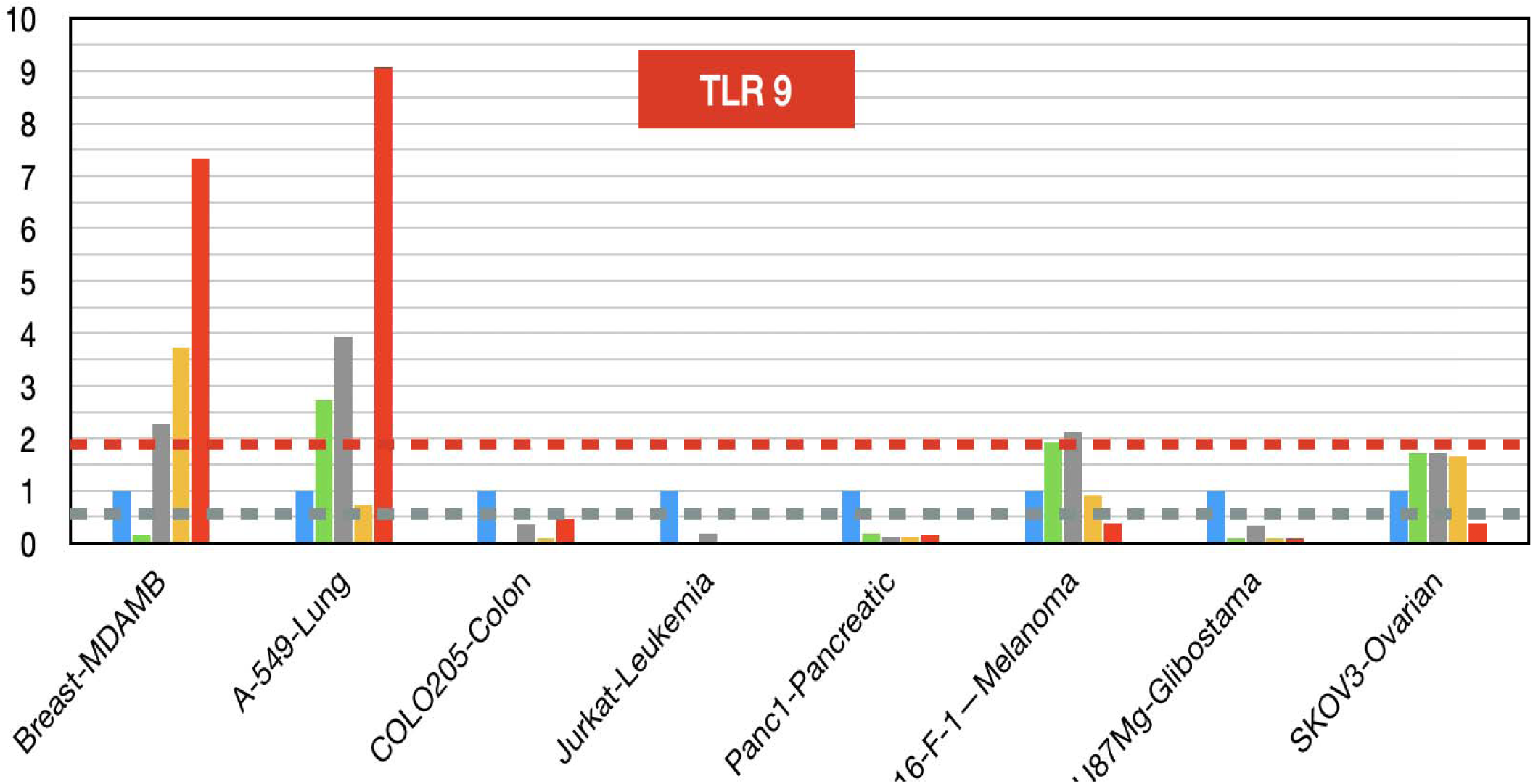

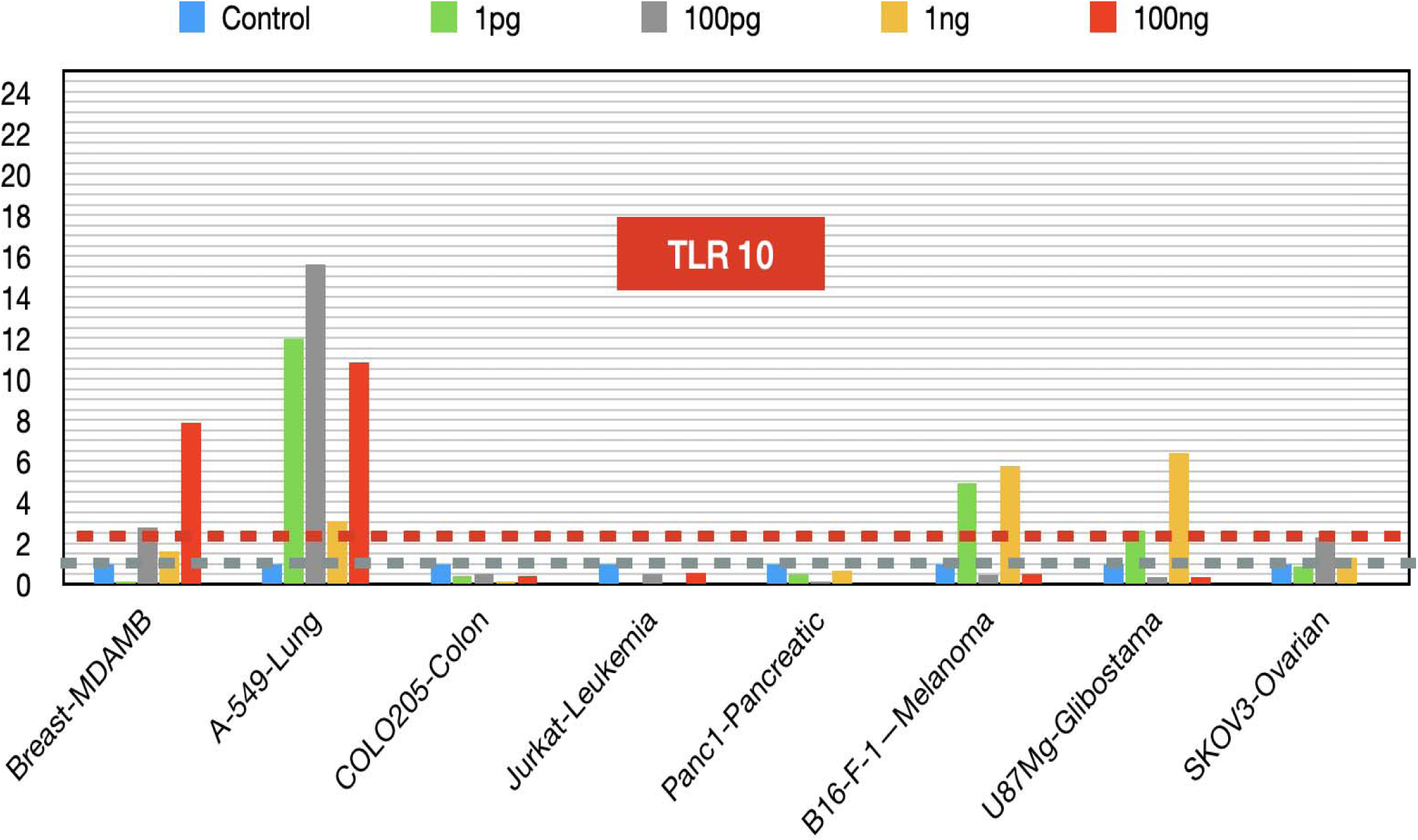

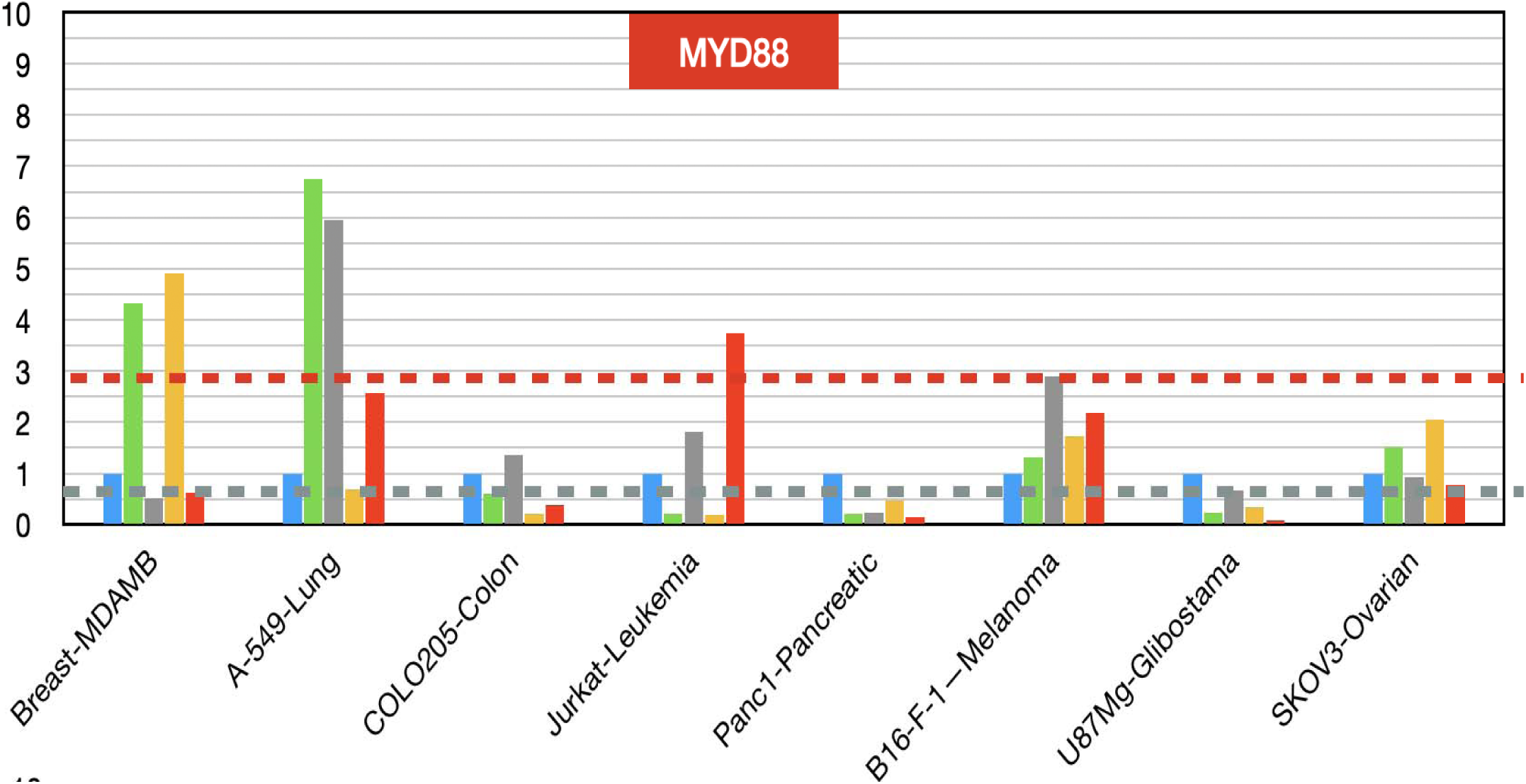

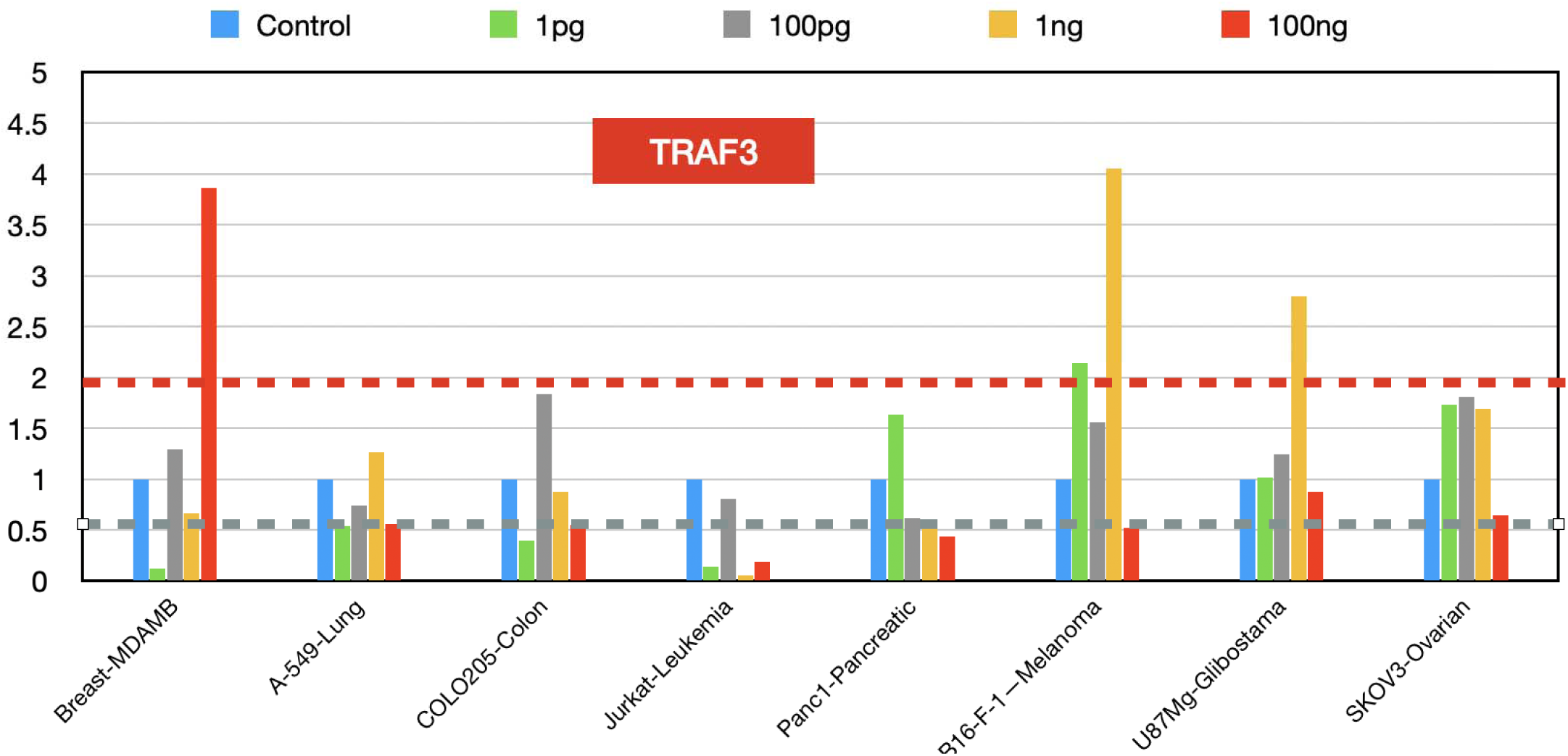

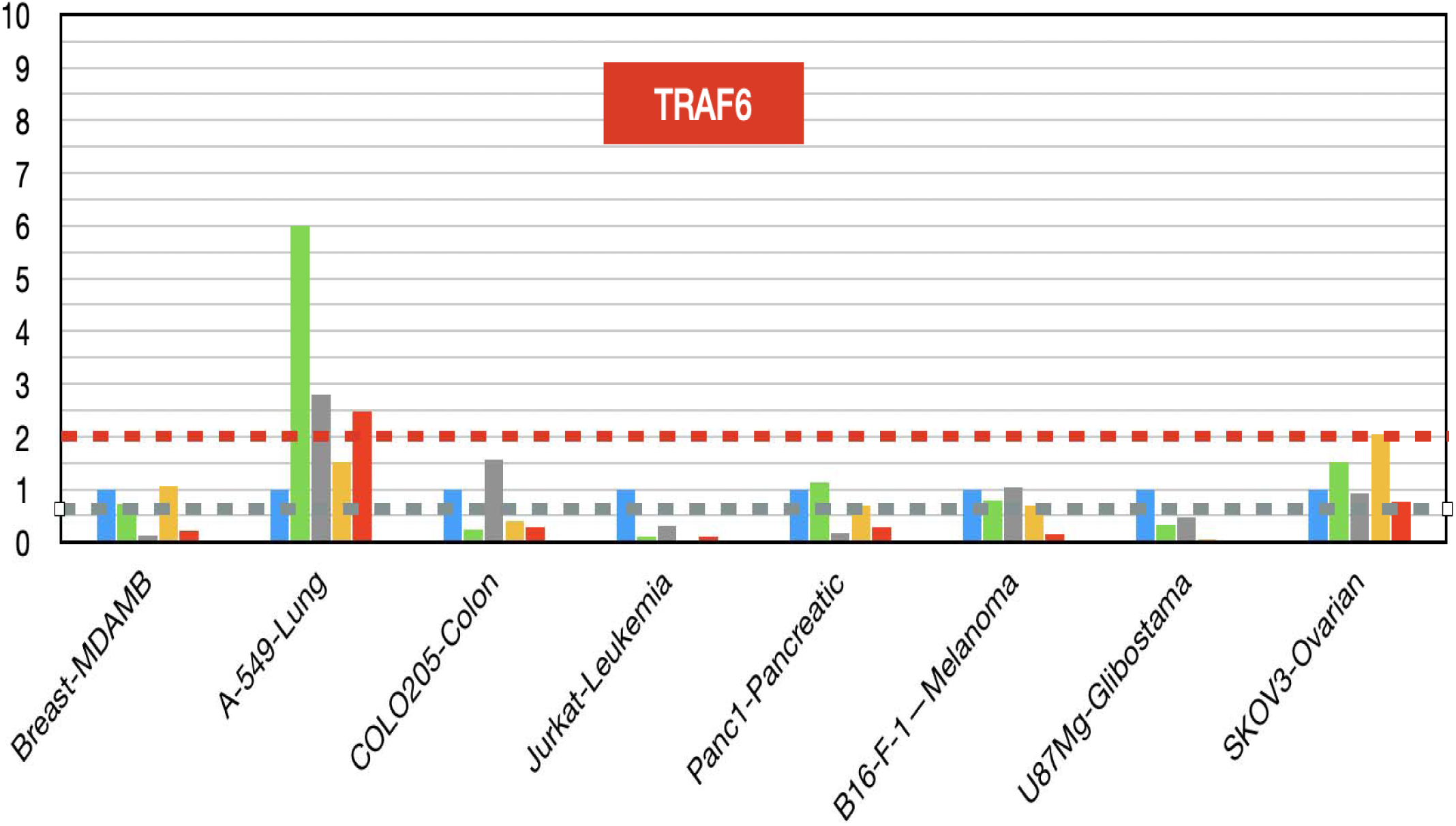

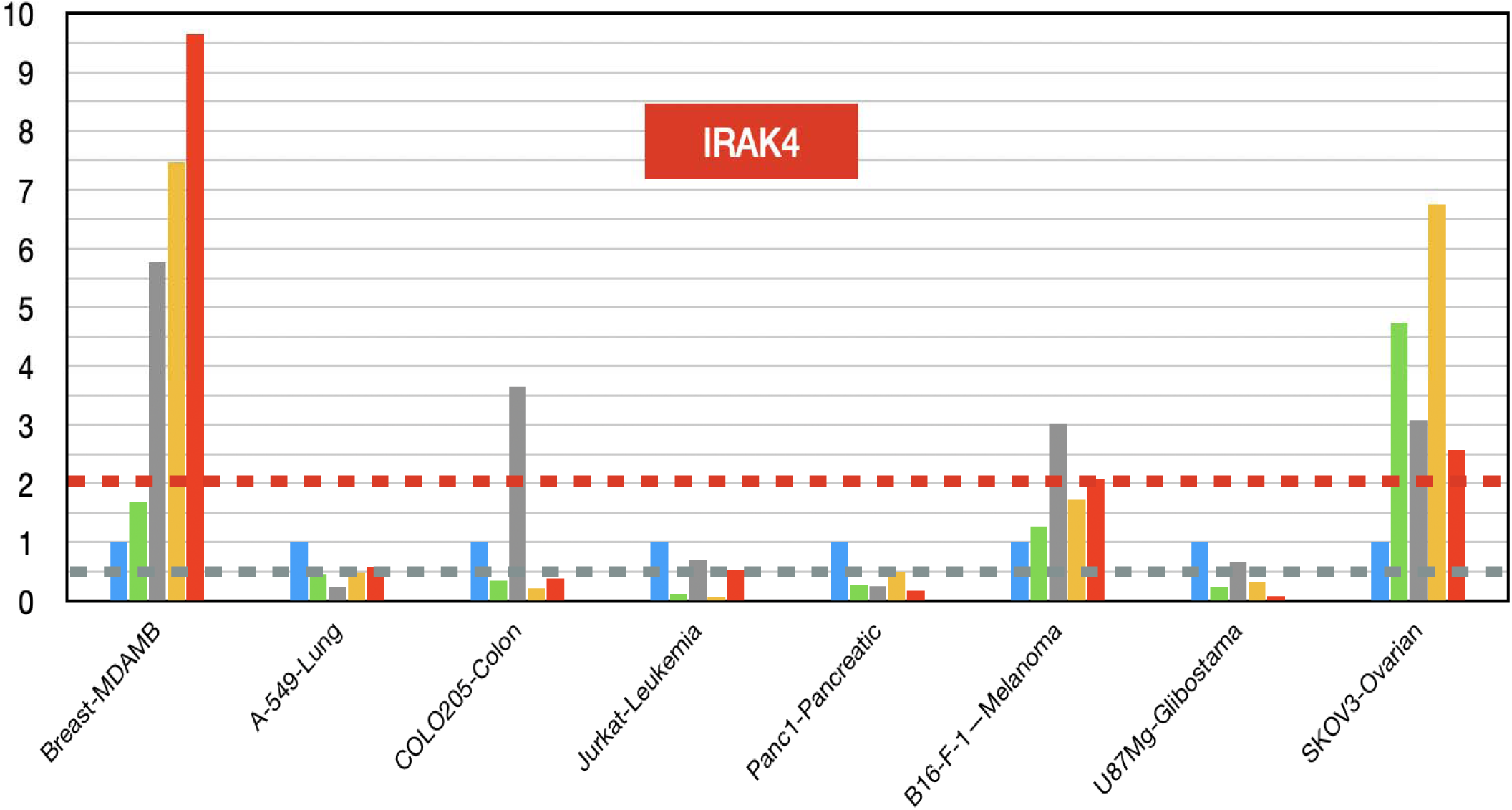

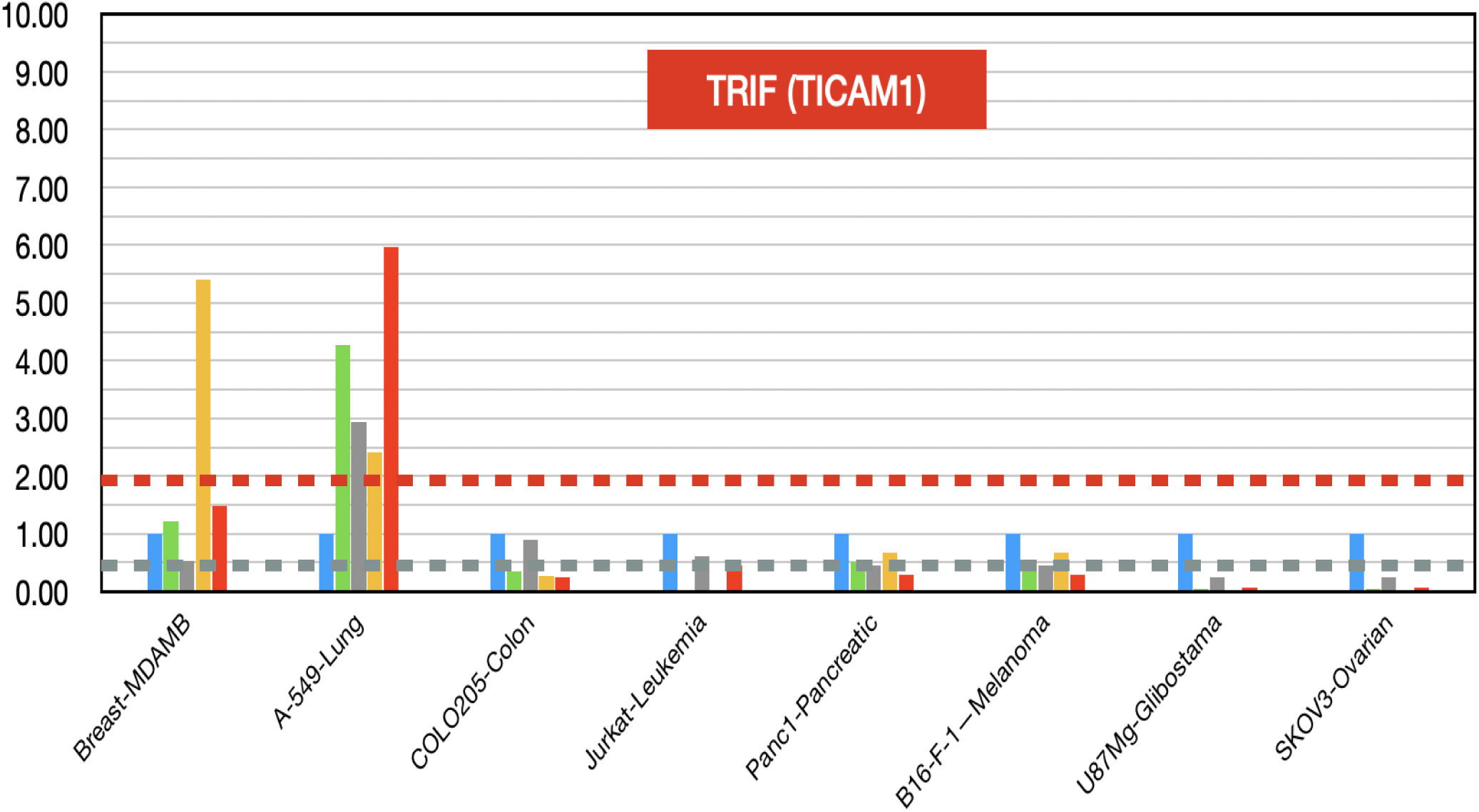

## Discussion

The expression of TLRs in cancer cells is linked to various outcomes, including resistance to chemotherapy and tumor development(6).

TLRs have been identified as potential marker molecules for developing neoplastic diseases and are increasingly highlighted as prognostic factors in various cancers (7). Additionally, the diverse responses of cancer cells from the exact histological origin to the same TLR ligand indicate that the initial expression levels of TLRs can vary significantly among different cancer cell lines(8). This variability highlights the complexity of TLR expression and its implications for cancer therapy.

In our previous work (2) on the effect of metadichol on TLR expression in PBMCs, we discussed in detail how TLR expression is connected to various nuclear receptors and sirtuins, which are also expressed by metadichol in somatic cells. This paper discusses the effects of metadichol expression on TLR expression in cancer cells and downfield genes in the TLR network and their implications for cancer.

### TLR1

Inhibition of TLR1/2 signaling reduces the response to chemotherapy and leads to fewer tumor-infiltrating immune cells(9).

### TLR2

Targeting Tlr2 reduces lung tumor growth in mouse models. Its inhibition enhances chemotherapy efficacy (10). It is associated with improved survival and premalignant regression in NSCLC (11) and mediates oncogene-induced senescence and the SASP (senescence-associated secretory phenotype) in early lung tumors (12). The Tlr2-mediated SASP recruits myeloid cells and promotes senescence surveillance.

### TLR3

The role of TLR3 in activating immune responses means that its absence can lead to diminished immune attack on tumors (13). Increased expression leads to tumor regression and increased carcinogenesis and has potential applications in resistant cancer.

### TLR 4

The reduced expression of TLR4 across cancer lines (14) indicates reduced inflammation, and increased TLR4 expression in non-small cell lung carcinoma (NSCLC) cells is an anticancer agent, indicating that TLR4 activation could be beneficial in specific cancer types (15,16).

### TLR5

High expression has been associated with better prognosis and antitumor effects in colorectal and prostate cancers. The activation of TLR5 by flagellin (17) inhibited increased breast cancer cell proliferation and colony formation in vitro and stimulated antitumor immunity by activating innate immune cells and promoting CD8+T-cell activation and tumor infiltration. These findings suggest that TLR5 agonists could be explored as potential therapeutic agents, particularly in combination with other immunotherapies(18).

### TLR6

It forms heterodimers with TLR2 and is involved in recognizing bacterial lipopeptides. The role of TLR6 in cancer prognosis appears to be context-dependent: In some cancers, such as colorectal cancer (19), decreased TLR6 expression is associated with cancer progression, suggesting a potential protective role of TLR6 against tumor development. In contrast, high TLR6 expression in esophageal squamous cell carcinoma (20) is linked to improved prognosis, indicating that TLR6 might have an antitumorigenic effect in this context. Overall, TLR6 expression levels can serve as a biomarker for cancer prognosis, but their role can vary depending on the type of cancer and the specific tumor microenvironment.

### TLR7

It is highly expressed in primary human ductal pancreatic cancer and is associated with advanced tumor stages and poor clinical outcomes in non-small cell lung carcinoma (NSCLC). Given the protumorigenic role of TLR7 in many cancers, TLR7 inhibitors (21) could be promising therapeutic strategies to reduce tumor growth, metastasis, and chemoresistance.

### TLR8

Its inhibitors have shown promise in reducing tumor growth (22) and enhancing immune responses in pancreatic cancer cell lines. TLR8 agonists enhance tumor immunity by preventing tumor-induced T-cell senescence, suggesting that TLR8 agonists could improve natural killer (NK) cell functions in cancer immunotherapy(23).

### TLR9

It plays a vital role in generating antitumor immune responses after chemotherapy. TLR9 agonists (24), such as CpG oligonucleotides, have shown promise in inhibiting tumor growth in neuroblastoma and other cancers. However, in cases where TLR9 activation promotes tumor progression, TLR9 inhibitors (25) could be promising therapeutic strategies. TLR9 has both anti-inflammatory and potential antitumor effects.

### TLR10

Agonists of TLR10 can help modulate the immune response and reduce tumor-promoting inflammation (26), whereas TLR10 inhibitors can bring out a reduction of tumor growth and metastasis in cancers where high TLR10 expression is correlated with poor outcomes (27,28).

### MYD88

MyD88) is an adaptor protein involved in the signaling pathways of Toll-like receptors (TLRs) and interleukin-1 receptors (IL-1Rs), which play crucial roles in the immune response and inflammation. Its expression and role in cancer have been extensively studied, revealing its involvement in tumor progression and immune response modulation and its potential as a prognostic marker and therapeutic target(31).

MyD88 expression and role in cancer. (29)

- **Colorectal cancer (CRC):** MyD88 is highly expressed in CRC tissues, particularly in patients with liver metastasis, and is associated with poor prognosis (30). High MyD88 expression is correlated with increased tumor growth and invasion and poor survival outcomes in CRC patients.
- In breast cancer, elevated MyD88 expression (31) is linked to aggressive tumor characteristics, including metastasis, recurrence, and drug resistance. It modulates inflammatory and chemotactic factors and influences the tumor immune microenvironment, making it a valuable prognostic marker and therapeutic target.
- **Ovarian cancer:** MD88 expression (32) has a dual role in ovarian cancer. In high-grade serous ovarian cancer (HGSOC), high MyD88 levels tend to be associated with advanced disease stages and shorter overall survival. Conversely, in low-grade serous ovarian cancer (LGSOC), high MyD88 expression is linked to better survival outcomes.
- **Diffuse large B-cell lymphoma (DLBCL):** A specific mutation in MyD88, L265P, is prevalent in DLBCL and Waldenström’s macroglobulinemia and promotes tumor cell survival and proliferation through NF-κB signaling (33).
- **Tumors:** MyD88 promotes tumor growth, invasion, and metastasis across various cancer types. It activates signaling pathways, such as the NF-κB and MAPK pathways, crucial for tumor cell proliferation and survival (34).
- **Tumor-Initiating Cells (TICs):** MyD88 signaling can generate TICs (35), particularly in p53-deficient cells, through the NF-κB-HIF-1α activation cascade. This mechanism is essential for inflammation-induced cancer development.
- **Immune Modulation:** MyD88 plays a role in immune escape by regulating the tumor microenvironment. It influences the recruitment and function of immune cells, such as macrophages, which can suppress antitumor immune responses (36).

MyD88 expression (37) serves as a potential biomarker for cancer prognosis, with its levels correlating with disease severity and patient outcomes. Targeting MyD88 and its signaling pathways offers a promising approach to cancer therapy, particularly in cancers where MyD88 contributes to tumor progression and immune evasion. Inhibitors of MyD88 or its downstream pathways, such as immune checkpoint inhibitors, could increase the efficacy of existing treatments.

### TRAF3

The dual role of TRAF3 as a tumor suppressor and immune modulator presents (38) opportunities for therapeutic intervention. Targeting TRAF3-related pathways could enhance antitumor immunity and inhibit tumor growth (39). For example, manipulating TRAF3 expression or function could improve the efficacy of existing cancer therapies, such as immune checkpoint inhibitors, by modulating the tumor microenvironment and immune response (40). In summary, TRAF3 is a critical regulator in cancer biology (41), and its functions vary across different cancer types and cellular contexts. Its role as a tumor suppressor and immune modulator makes it a promising target for cancer therapy.

### Traf6

TRAF6 plays a crucial role in cancer development and progression by promoting tumor growth, metastasis, chemoresistance, and immune evasion. Targeting TRAF6 offers a promising strategy for cancer therapy, particularly in cancers where TRAF6 is overexpressed and contributes to poor clinical outcomes(42–46).

### IRAK4

IRAK4 plays a significant role in cancer biology by promoting tumor progression, chemoresistance, and immune evasion. Its expression levels and activity across various cancers make it an attractive target, and ongoing research is focused on developing effective IRAK4 inhibitors for clinical use (48–49).

### TICAM1

It plays a multifaceted role in cancer by influencing immune responses, cell death pathways, and tumor progression. Its expression levels and functional roles vary across different cancers, making it a potential target for therapeutic intervention and a biomarker for cancer prognosis. (50–53)

The results show that metadichol regulates the expression of all ten Toll-like receptors (TLRs 1--10), as well as key signaling molecules such as MYD88, IRAK4, TICAM1 (TRIF), TRAF3, and TRAF6, in various cancer cell lines. This ability to modulate multiple components of the TLR signaling pathways simultaneously and in a concentration-dependent manner has significant implications for cancer treatment. These include immune modulation, targeting multiple pathways, dose-dependent effects, potential for combination therapy, and safety.

#### Immune Modulation

By influencing the expression of TLRs and related signaling molecules, metadichol can modulate the immune response. TLRs play crucial roles in recognizing pathogens and initiating immune responses. In cancer, they can either promote or inhibit tumor growth depending on the context. Metadichol ability to upregulate or downregulate these receptors and adaptors could be harnessed to enhance antitumor immunity or reduce tumor-promoting inflammation.

#### Targeting Multiple Pathways

Cancer is a complex disease often driven by multiple signaling pathways. The Metadichol multitarget approach, which affects several TLRs and downstream signaling molecules, aligns with the need for therapies to address the multifaceted nature of cancer biology. This approach could overcome the limitations of single-target therapies, which may not be effective because of cancer adaptability and redundancy in signaling pathways.

#### Dose-dependent effects

This characteristic could be leveraged to fine-tune the immune response in cancer therapy, optimizing the balance between stimulating an effective immune response and avoiding excessive inflammation that could promote tumor growth.

#### Potential for Combination Therapy

The ability of metadichol to modulate key signaling molecules, such as MYD88, IRAK4, and TRAF6, which are involved in pathways such as the NF-κB and MAPK pathways, suggests potential synergy with existing cancer treatments. For example, combining metadichol with immune checkpoint inhibitors or chemotherapy could enhance therapeutic efficacy by simultaneously targeting multiple aspects of tumor biology and the immune microenvironment.

#### Safety and versatility

Metadichol’s reported nontoxic profile (54–56) and ability to modulate a wide range of targets make it a promising candidate for further investigation in cancer therapy. Its versatility in affecting various signaling pathways suggests it could be used to treat various cancers.

##### Summary

The downregulation of Toll-like receptors (TLRs) at high concentrations in cancer cell lines can be a significant antitumor effect. TLRs are involved in the immune system by recognizing pathogens and activating immune responses. Downregulating TLRs can reduce chronic inflammation, often associated with tumor progression. (57). This modulation can create a less favorable environment for cancer cells. Chronic inflammation can promote tumor growth and survival. The inflammatory signals that support tumor growth can be diminished by downregulating TLRs, potentially slowing down or inhibiting cancer progression (58).

TLR downregulation can also enhance the effectiveness of other cancer therapies, such as immunotherapy. The immune system can be targeted by reducing the suppressive signals in the tumor microenvironment (59). Some studies suggest that altering TLR signaling (60) results in programmed cell death (apoptosis) in cancer cells, further contributing to the antitumor effect. Also, specific TLRs can be upregulated in certain cancers, as in TLR4 in lung cancer cells. This ability of metadichol to modulate the expression of TLRs and downfield transcription factors in the right direction represents a novel approach for cancer treatment, offering potential benefits in terms of immune modulation, pathway targeting, and combination therapy.

## Supporting information

raw Data

## Supplementary Data

Access to data and materials

The manuscript and supplementary resources contain all the raw data.

## Conflicting goals

The author is the founder and principal shareholder of Nanorx, Inc., NY, USA.

## Funding

This study was funded internally by Nanorx, Inc., NY, USA.

## Supplementary files

**Name of File.** Q-RT-PCR Raw Data:

This file contains raw data on fold changes by qRT-PCR and an evaluation of the effects of metadichol on the gene expression of Toll-like receptors on cancer lines.

## Notes

### Competing Interest Statement

The author is founder and CEO of Nanorx Inc in which he is a major share holder.

### Summary of Updates

Title changed to reflect correct information about the paper

